# Nodal Modulator is required to sustain endoplasmic reticulum morphology

**DOI:** 10.1101/2021.01.28.428001

**Authors:** Catherine Amaya, Christopher JF Cameron, Swapnil C. Devarkar, Mark B. Gerstein, Yong Xiong, Christian Schlieker

## Abstract

Nodal Modulator (NOMO) is a widely conserved type I transmembrane protein of unknown function, with three nearly identical orthologs specified in the human genome. We identified NOMO1 in a proteomics approach aimed at the identification of proteins that support the structural integrity of the endoplasmic reticulum (ER). Overexpression of NOMO1 imposes a sheet morphology on the ER, while depletion of NOMO1 and its orthologs causes a collapse of ER morphology concomitant with the formation of membrane-delineated holes in the ER network. These structures are positive for the autophagy marker LAMP1, and LC3 is profoundly upregulated upon NOMO depletion. *In vitro* reconstitution of NOMO1 revealed a dimeric state that is mediated by the cytosolic tail domain, with each monomer featuring a “beads on a string” structure likely representing bacterial Ig-like folds. Based on these observations and a genetic epistasis analysis including the known ER-shaping proteins Atlastin2 and Climp63, we propose a role for NOMO1 in the functional network of ER-shaping proteins.

## Introduction

As the largest, single-membrane bound organelle, the endoplasmic reticulum (ER) is responsible for critical and diverse functions, including lipid synthesis, folding and export of membrane and secretory proteins, and calcium storage (Ma & Hendershot, 2001; Matlack, Mothes, & Rapoport, 1998; Meldolesi & Pozzan, 1998). These responsibilities are divided into three structurally distinct regions, namely the nuclear envelope (NE), sheets, and tubules (Palade, 1956). These regions partition protein synthesis and folding to the sheets, and organelle fission and calcium storage to tubules (Friedman et al., 2011). The structural integrity of these regions is maintained and regulated by unique membrane shaping proteins.

The membrane shaping proteins necessary to support the curvature of ER tubules have largely been established (Powers, Wang, Liu, & Rapoport, 2017), which include Reticulons (RTNs), Atlastins (ATLs), and receptor expression enhancing proteins (REEPs) (Hu et al., 2009; Voeltz, Prinz, Shibata, Rist, & Rapoport, 2006). The prominent structural motif shared by these proteins is a transmembrane hairpin, which serves as a wedge that is inserted into the outer lipid layer of the ER membrane to impose high curvature on the membrane and help create a tubular shape. Additionally, ATLs have a cytosolic GTPase domain responsible for fusion and tethering of tubules, and creating the connected reticular network of the ER (Hu et al., 2009). Depletion of ATLs results in ER tubules becoming abnormally long and unbranched and disrupts ER tubule functionality (Rismanchi, Soderblom, Stadler, Zhu, & Blackstone, 2008; G. Zhao et al., 2016). This disruption demonstrates the critical role of maintaining ER membrane morphology for the function of the ER. The importance of understanding how the ER maintains structural integrity is highlighted by diseases that occur when the functions of ER shaping proteins are disrupted. Mutations in tubule shaping proteins, such as in ATLs, spastin, RTNs, and REEP1 are associated with diseases such as amyotrophic lateral sclerosis, hereditary spastic paraplegia (HSP), and other neurodegenerative disorders (Blackstone, O’Kane, & Reid, 2011; Chiurchiu, Maccarrone, & Orlacchio, 2014; Park, Zhu, Parker, & Blackstone, 2010).

Although tubule shaping proteins have been well established, much remains to be learned about sheet morphology. The maintenance of sheet spacing is largely attributed to Climp63, an ER resident-microtubule binding protein that features a long coiled-coil domain in the ER lumen (Klopfenstein, Kappeler, & Hauri, 1998; Shibata et al., 2010; Vedrenne, Klopfenstein, & Hauri, 2005), whereas the high curvature edges of the sheets are stabilized by tubule shaping proteins like RTNSs (Jozsef et al., 2014; Schroeder et al., 2019; Voeltz et al., 2006). Initially, it was proposed that the coiled-coil domain of Climp63 dimerizes across the ER lumen to support an intermembrane distance of about 60 nm (Shibata et al., 2010). Indeed, modulating the length of the Climp63 coiled-coil domain was shown to correlatively affect the ER luminal distance (B. Shen et al., 2019). Kinectin and p180 have also been proposed to contribute to the flatness of sheets. Despite these contributions to maintaining sheet morphology, simultaneous depletion of Kinectin, p180, and Climp63 or Climp63 alone does not result in a loss of sheets. Rather, the ER diameter is uniformly decreased to 30 nm (B. Shen et al., 2019; Shibata et al., 2010). Climp63 has also been proposed to keep the opposing sheet membranes from collapsing into each other (Schweitzer, Shemesh, & Kozlov, 2015). However, Climp63 depletion does not lead to a loss of sheets, and no functional perturbations of the ER have been reported. These observations suggest that additional, yet unidentified, sheet shaping proteins exist to support sheet formation and prevent disruption to ER sheet functions.

Here, we use a proximity ligation-based approach to identify additional ER-luminal proteins that could contribute to membrane spacing. We identified Nodal Modulator 1 (NOMO1), a widely conserved type1 transmembrane glycoprotein, as an abundant luminal constituent of the ER. Depletion of NOMO1 in a tissue culture model perturbs ER morphology, while its overexpression imposes a defined intermembrane spacing on the ER. Furthermore, *in vitro* reconstitution including light scattering and low-resolution electron microscopy (EM) collectively suggest that NOMO1 is a parallel dimer of rod-shaped molecules, featuring Ig-folds that are arranged as “pearls on a string”. Based on these observations, as well as a genetic epistasis analysis including several ER-shaping proteins, we place NOMO1 in a functional network of proteins responsible for establishing and maintaining the morphology of the ER.

## Results

### Identification of NOMO1 as an abundant, ER-luminal protein

To identify potential sheet shaping proteins, we employed a proximity ligation approach. Previous proteomes of the ER were obtained by subcellular fractionation-based techniques that encompassed the entire ER membrane network (Chen, Karnovsky, Sans, Andrews, & Williams, 2010; Sakai, Hamanaka, Yuki, & Watanabe, 2009), whereas we were specifically interested in the ER lumen. To this end, we used an engineered monomeric peroxidase (APEX2) (Lam et al., 2015). In the presence of hydrogen peroxide, APEX2 creates biotin-phenoxyl radicals that will biotinylate proteins in a 20 nm radius (Hung et al., 2016; Hung et al., 2014; Rhee et al., 2013). We employed ER-APEX2, a construct previously shown to specifically localize to the ER lumen by virtue of a signal sequence (Lee et al., 2016). This construct was expressed in HeLa cells that were then incubated with biotin and treated with hydrogen peroxide to conjugate biotin to ER luminal proteins. The control sample was transfected with ER-APEX2 but no hydrogen peroxide was added. The treated cells were lysed in SDS buffer, and a streptavidin bead resin was used to isolate biotinylated proteins. To control for labeling efficacy, samples were eluted and subjected to SDS-PAGE and blotting using a streptavidin conjugate for detection. Since robust, hydrogen-peroxide dependent labeling was observed for a variety of proteins (Fig. 1A), we performed an analogous experiment on a larger scale and analyzed the resulting eluates via mass spectrometry following tryptic digestion. As expected, the most abundant species identified included constituents of ER protein synthesis and folding machinery (Fig. 1B), including the ER chaperones BiP, PDI, Endoplasmin and CCD47, all of which are known residents of the ER lumen (Chitwood & Hegde, 2020; Helenius & Aebi, 2004). In addition, NOMO2 and NOMO1 were the eighth and ninth most abundant proteins identified as judged by spectral counts, with high sequence coverage (48%) (Fig. 1B).

**Figure 1.**
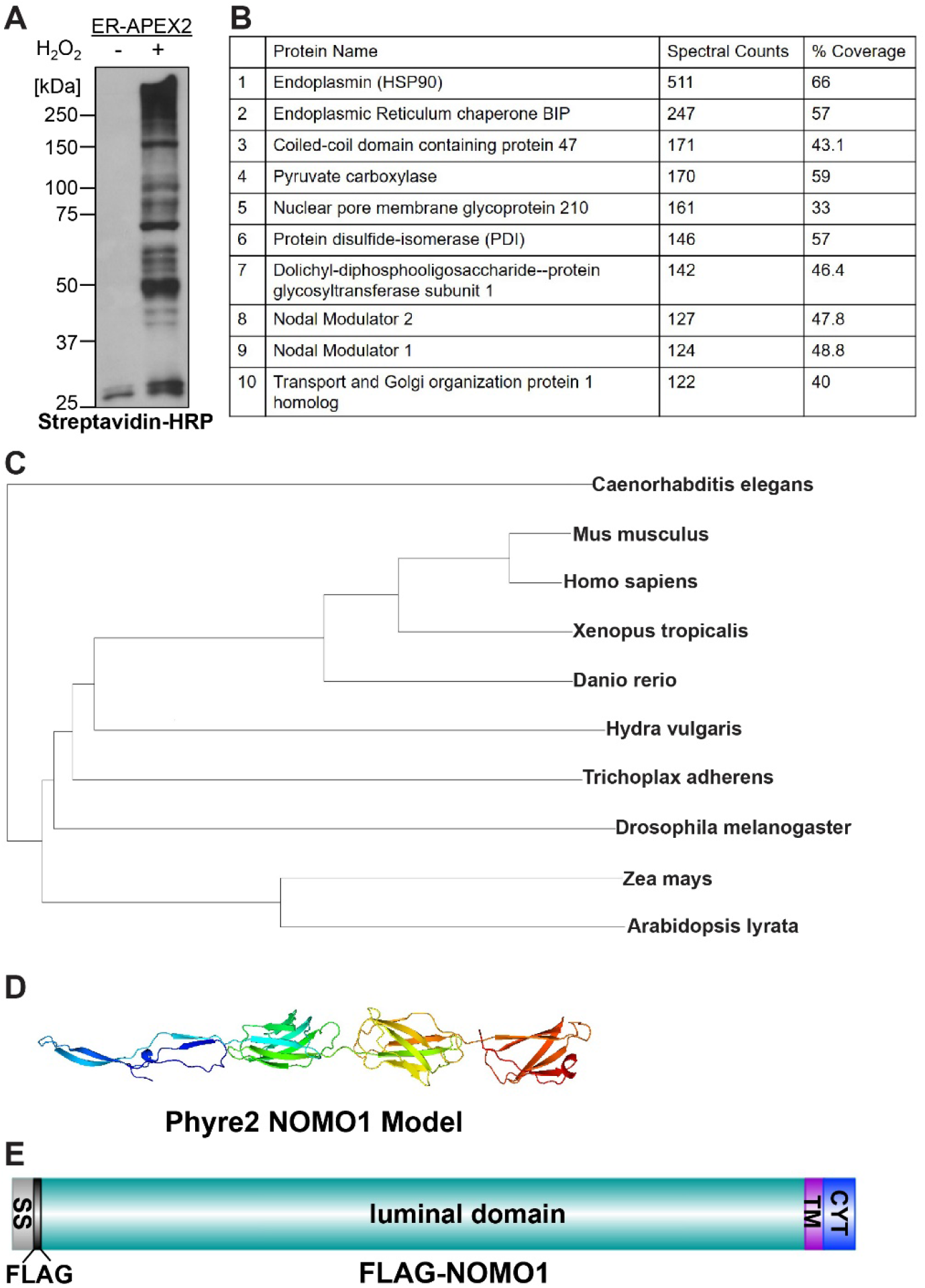
Identification of NOMO1 as abundant and conserved ER-resident protein. A. Cells expressing ER-APEX2 were treated with biotin-phenol in absence or presence of hydrogen peroxide, lysed and subjected to western blotting using streptavidin-HRP. B. Table of top ten most abundant proteins from mass spectrometry analysis in order of spectral count; % Coverage is the sequence coverage of the protein based on the peptide sequences identified. C. Phylogenetic tree of NOMO1 homologs in indicated metazoan and plant species. D. FLAG-NOMO1 domain structure. Note that the FLAG tag was inserted between the cleavable signal sequence (SS) and the luminal domain. TM, transmembrane domain, CYT, cytosolic tail.

NOMO1 is a type I transmembrane protein that is conserved across all metazoans (Haffner et al., 2004). Notably, NOMO homologs are also present in plants, both in monocotyledones (*Zea mays*) and dicotyledons (*Arabidopsis lyrata*) (Fig. 1C). While other metazoan organisms specify a single copy of NOMO, three copies of NOMO are present in the human genome designated: *NOMO1*, *NOMO2*, and *NOMO3* (Yates et al., 2019). NOMO1 and NOMO2 specify a 134 kDa membrane protein composed of an N-terminal 1124 residue luminal domain, a transmembrane domain, and a short, 40 residue cytosolic domain. The luminal domains of the three proteins are identical except for six amino acids (Fig. S1A). NOMO2 has a cytosolic domain that is 45 residues longer than NOMO1 and NOMO3, resulting in a 139 kDa membrane protein. This extremely high similarity suggests that NOMO orthologs have arisen from recent gene duplication events and have identical or similar cellular functions.

To begin to understand which function NOMO might have in the ER, we employed BLAST searches, secondary structure predictions, and fold recognition programs to identify homology to proteins of known structure. While these searches did not reveal related human proteins, NOMO1 is predicted to form a beta sheet-rich structure (Fig. S1B) by PSIPred (Buchan & Jones, 2019). Consistently, a significant structural degree of similarity was detected between NOMO1 and several bacterial Ig-like fold proteins. The highest similarity was observed for BaTIE, a sortase-anchored surface protein from *Bacillus anthracis* (Miller, Banfield, & Schwarz-Linek, 2018), featuring 4 tandem Ig domains of 19 nm in length (Fig. S1C). Phyre2 (Kelley, Mezulis, Yates, Wass, & Sternberg, 2015) modeled NOMO1 residues 58-398 with 99% confidence (Fig. S1D), predicting 4 consecutive Ig folds for this region (Fig. 1D). This structural homology led us to hypothesize that NOMO1 might adopt an extended rod structure that could serve as a structural component to support membrane spacing.

### NOMO depletion results in altered ER morphology

As a first test to determine if NOMO depletion contributes to ER morphology, we depleted NOMO in U2OS cells using siRNA. Due to the high genomic similarity between *NOMO1, NOMO2*, and *NOMO3*, siNOMO1 targets all three corresponding mRNAs. In the following, we will refer to the experimental condition simultaneously depleting NOMO1, NOMO2, and NOMO3 as NOMO. The canonical nomenclature of NOMO1 will be used for experiments based on the specific NOMO1 cDNA or protein. NOMO depletion caused a striking rearrangement of the ER network and large holes in the ER of up to 5 μm in diameter were visible by immunofluorescence microscopy (Fig. 2A). Attempts at generating a CRISPR/Cas9 NOMO KO cell line were unsuccessful. While single cell colonies were obtained in which the hole phenotype was visible, cells were not viable in culture after several passages, suggesting an important, if not essential function.

**Figure 2.**
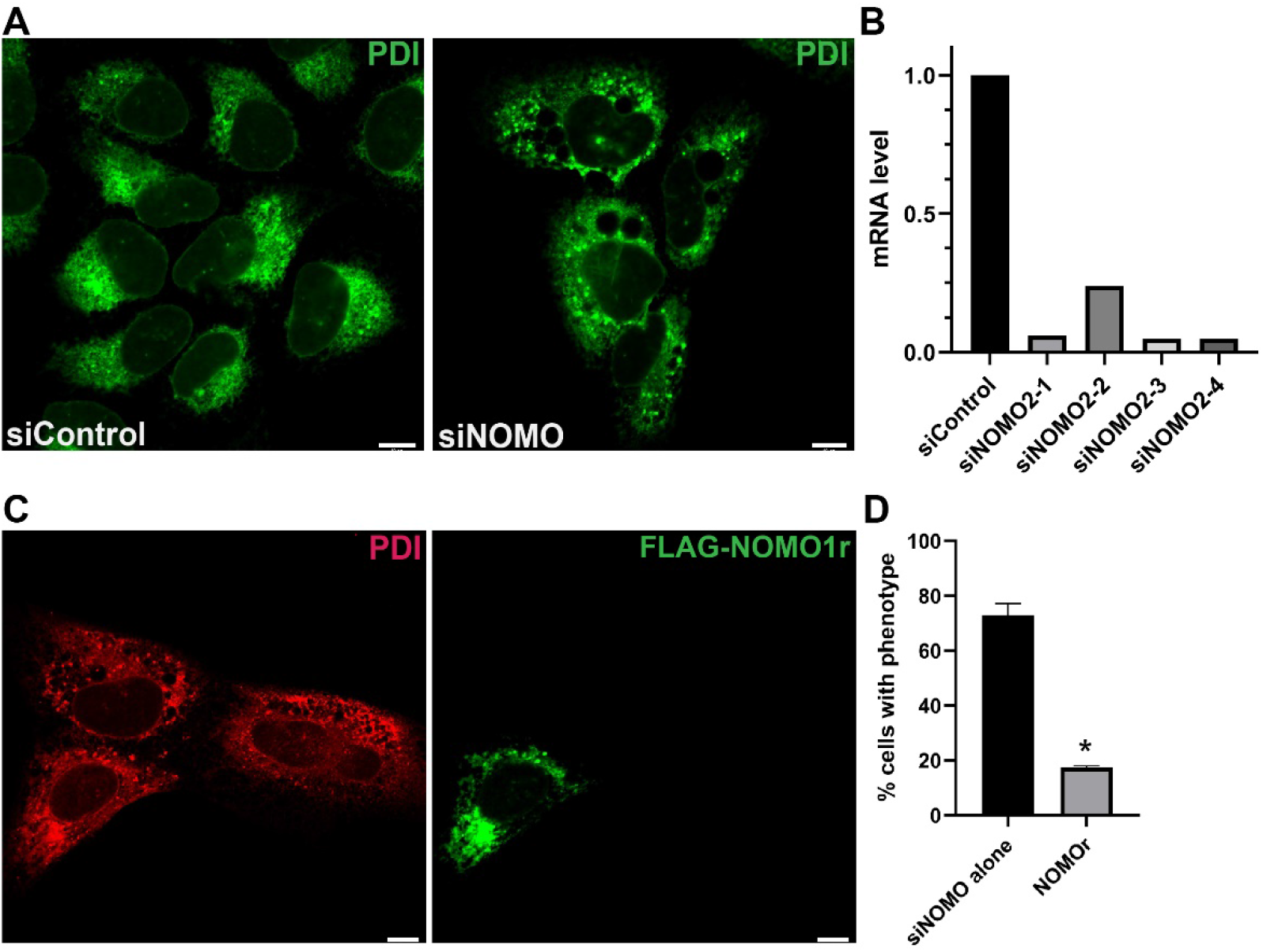
NOMO depletion results in profound changes of ER morphology. A. U2OS cells were transfected with the respective siRNA for 48 hours. B. Quantification of mRNA level of each NOMO siRNA by qPCR. C. Representative image of phenotypic rescue of the NOMO knockdown phenotype by an siRNA-resistant construct, FLAG-NOMO1r. D. Quantification of rescuing ability of FLAG-NOMO1r, n=100, N=3, p<0.05. Asterisks denote P<0.005 compared to control. Error bars indicate standard deviation. All scale bars are 10 μm.

To demonstrate that the siRNA-induced phenotype was specifically due to NOMO depletion, a NOMO1 rescue construct, FLAG-NOMO1r, was designed by introducing silent mutations into the targeting site of siRNA #3. This siRNA depleted NOMO mRNA by over 90% as quantified by qPCR (Fig. 2B). FLAG-NOMO1r reproducibly reduced the ER phenotype from 68% penetrance to 20%, providing further evidence that the hole phenotype observed is specifically caused by NOMO depletion (Fig. 2C, D). Since the simultaneous depletion of all three NOMO orthologs can be rescued by FLAG-NOMO1r alone, we conclude that NOMO1 has a major function in the context of ER morphology.

### Genetic interactions between NOMO and known ER-shaping proteins

From a topological perspective, the predicted domain architecture of NOMO is reminiscent of the structural domain composition of Climp63 that includes a sizeable luminal domain expanding into the ER lumen, a transmembrane domain, and a short cytosolic tail (Vedrenne et al., 2005). Therefore, we sought to compare whether Climp63 depletion caused similar defects in ER morphology as NOMO depletion. Depletion of Atl2 was included as a tubule shaping protein for comparison. Surprisingly, Atl2 depletion resulted in strikingly similar holes as those caused by NOMO depletion, while Climp63 depletion had no effect on ER morphology when visualized by immunofluorescence microscopy (Fig. 3A).

**Figure. 3.**
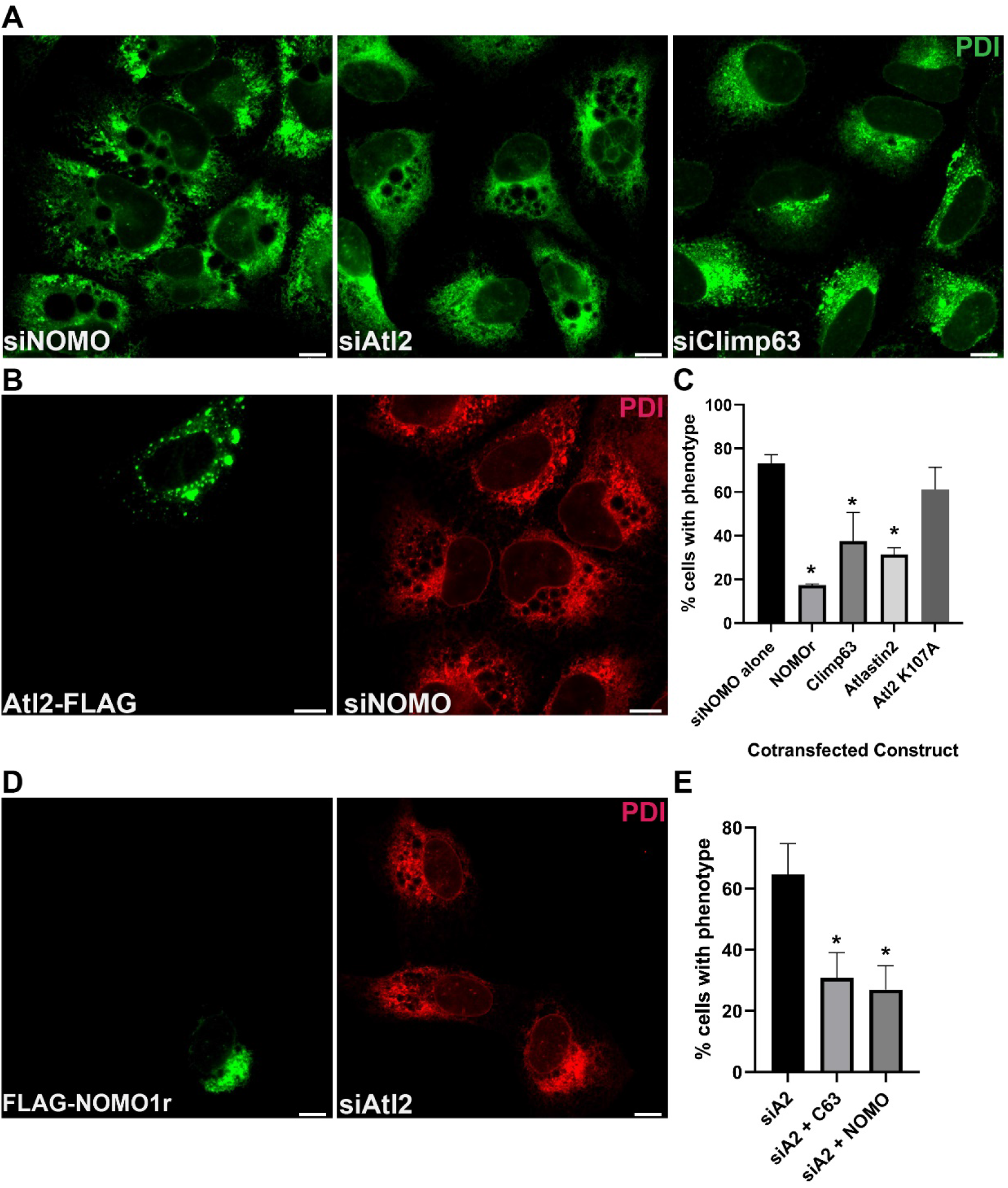
Epistasis analysis of known ER shaping proteins and NOMO1. A. U2OS cells were treated with respective siRNA for 48 hours and stained with PDI as an ER marker. B. Representative image of Atl2-FLAG overexpression (left panel) rescuing the NOMO KD phenotype as judged by PDI staining (right panel). C. Quantification of the ability of ER shaping proteins to rescue the NOMO KD ER phenotype, n=100, N=3. Error bars indicate standard deviation. D. Representative image of FLAG-NOMO1r overexpression (left panel) rescuing Atl2 KD phenotype (right panel). E. Quantification of NOMO1 and Climp63 overexpression rescuing the Atl2 KD phenotype, n=100, N=3. Error bars indicate standard deviation. Asterisks denote P<0.005. Scale bars are 10 μm.

Next, we asked if NOMO exhibits epistatic relationships with ATL2 or Climp63. First, we tested whether the overexpression of these known ER-shaping proteins modulates the observed hole phenotype. We transfected Atl2-FLAG into NOMO depleted cells and observed that Atl2-FLAG overexpression could significantly rescue the NOMO knockdown phenotype (Fig. 3B, C). Since Atl2 is required for ER fusion, we hypothesized that the fusogenic activity is required for this effect. To this end, a rescue assay was performed with a GTPase mutant of Atl2 that cannot fuse ER membranes, Atl2 K107A (Morin-Leisk et al., 2011). This Atl2 mutant did not rescue the NOMO knockdown hole phenotype (Fig. 3C), indicating that the rescue ability of Atl2 relies on the fusogenic activity. Furthermore, in an analogous experiment, we found that Climp63-FLAG did rescue the hole phenotype under NOMO depletion to a similar extent compared to Atl2 (Fig. 3C). These results suggest possible functional redundancy between NOMO and Climp63, assuming the hole phenotype is due to a lack of structural support for the sheets.

Lastly, since Atl2 depletion results in a similar hole phenotype, we performed the reciprocal rescue assays of co-transfecting NOMO1-FLAG or Climp63-FLAG into Atl2 depleted cells. We found that NOMO-FLAG and Climp63-FLAG both significantly reduced the penetrance of the Atl2 depletion phenotype (Fig. 3D, E). In conclusion, the observed genetic interactions among NOMO1, Climp63, and Atl2 are consistent with the interpretation that NOMO contributes to the elaborate network of ER-shaping proteins.

### Ultrastructural and compositional characterization of hole phenotype

To further explore the relationship between holes and the ER membrane, we processed U2OS cells depleted of NOMO for transmission electron microscopy (TEM). The holes were of significant size with an average area of about 1.2 um^2^ (Fig. 4A, C). Furthermore, holes often appeared to be devoid of any internal electron density and were delineated by membranes in various instances. In general, we encountered fixation issues resulting in suboptimal preservation of holes, possibly due to their large size and low interior content. While these fixation issue generally complicated direct visualization of membrane continuity, we observed in several cases that multiple membranes surrounded one hole (Fig. 4A, bottom panel). For comparison, we performed TEM analysis of U2OS cells under Atl2 depletion and observed similar membrane delineated holes (Fig. 4B). These results support the idea that a similar net result is obtained in response to the depletion of either NOMO or Atl2. Lastly, we noted electron-dense structures adjacent to or inside a subset of the holes under NOMO depletion (Fig. 4A, top and middle panels).

**Figure 4.**
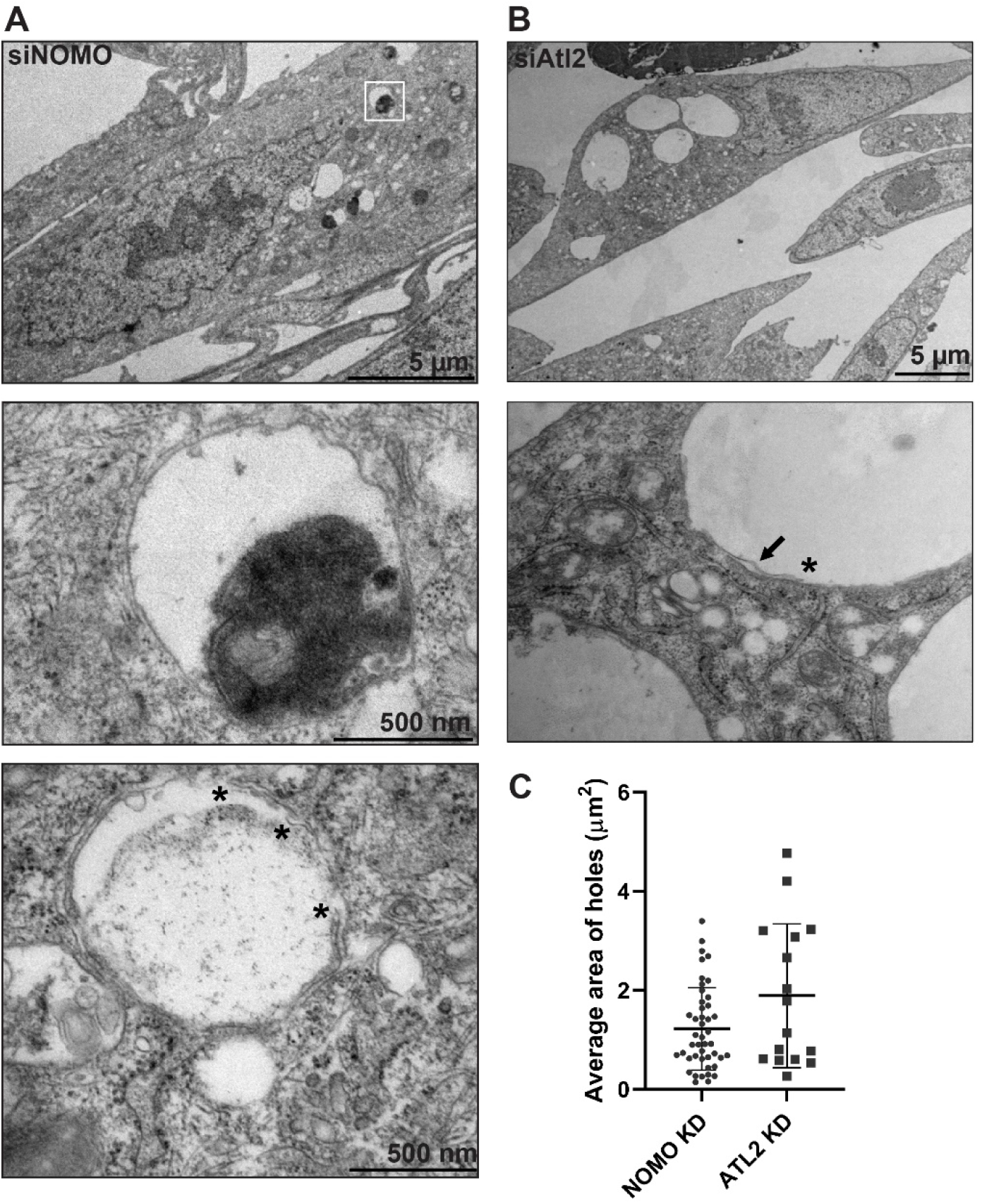
EM analysis of NOMO and Atl2-depleted cells. A. U2OS cells were successively treated with two doses of siNOMO 24 hrs apart and fixed 48 hrs after the second dose for EM processing. White square in the top panel identifies selection for middle panel. Asterisks in bottom panel denote free membrane ends. B. U2OS cells successively treated with two doses of siAtl2 as described in (A). Arrow in second panel indicates an identified membrane outlining the hole. C. Quantification of the area of observed gaps in A and B quantified on ImageJ. Error bars indicate standard deviation.

To determine if these electron-dense structures represent lysosomal compartments, U2OS cells were treated with siNOMO, siAtl2, or siClimp63 and analyzed by immunofluorescence microscopy using a lysosomal-associated protein 1 (LAMP1)-specific antibody. Indeed, we observed a large accumulation of LAMP1 signal in the ER holes resulting from NOMO and Atl2 depletion (Fig. 5A). The observed increase in lysosome size and accumulation compared to control cells could be an indicator of increased autophagy (de Araujo, Liebscher, Hess, & Huber, 2020). To address this point, we monitored LC3 processing by immunoblotting. LC3-I is processed to LC3-II as lysosomes increase their autophagic activity (Tanida, Ueno, & Kominami, 2008). We observed an increase of LC3-II under NOMO depletion compared to control, which is indicative of autophagy induction or dysregulation (Fig. 5B). We did not observe an increase in BiP levels under NOMO depletion, which would have indicated an induction of the unfolded protein responses (UPR) due to ER stress (Fig. 5B) (Walter & Ron, 2011). We also monitored LC3 processing under the depletion of ER shaping proteins Climp63 and Atl2, and NOMO binding partners TMEM147 and Nicalin (Dettmer et al., 2010; Haffner, Dettmer, Weiler, & Haass, 2007). An increase in LC3-II was also observed upon Climp63 depletion, though less pronounced compared to NOMO depletion (Fig. 5C), whereas the other tested conditions did not significantly increase LC3-II levels. Therefore, only Climp63 and NOMO depletions lead to an increase of LC3-II, indicative of autophagy induction or dysregulation.

**Figure 5.**
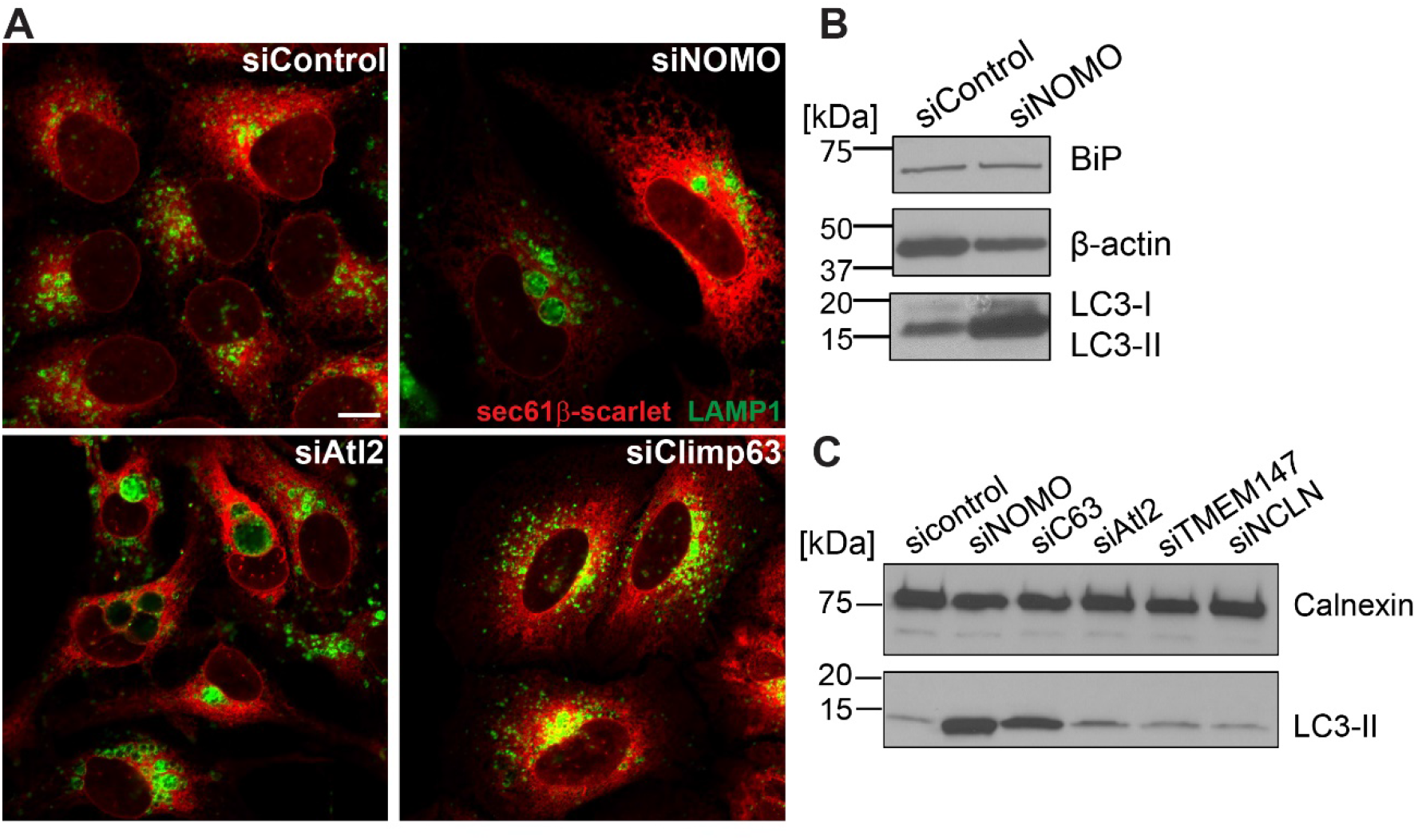
Sheet disruption increases autophagy. A. Representative images of U2OS cells treated with the respective siRNA to identify lysosome localization using LAMP1 as a marker. Scale bar is 10 μm. B. Immunoblot derived from control and NOMO depleted cells using the indicated antibodies. C. Immunoblot with calnexin and LC3 antibodies using U2OS cells extracts treated with the indicated siRNA.

### NOMO overexpression imposes ER sheet morphology

We hypothesized that if NOMO contributes to ER intermembrane spacing similar to Climp63, then overexpressing NOMO1 should affect the spacing of the ER lumen (B. Shen et al., 2019). To test this hypothesis, we overexpressed FLAG-NOMO1 in U2OS cells and imaged cells using PDI as the ER marker. We observed enlarged, continual ER areas reminiscent of sheets by confocal microscopy compared to the shorter structures of untransfected cells (Fig. 6A). To determine if ER sheet spacing was affected, we subjected HeLa cells overexpressing FLAG-NOMO1, as well as control cells transfected with empty vector, to TEM imaging. Cells overexpressing FLAG-NOMO1 had a constricted ER lumen diameter compared to control cells (Fig. 6C, D). When quantified, FLAG-NOMO1 overexpression reduced the diameter of the ER lumen from an average intermembrane distance of 80 nm to 30 nm (Fig. 6E). Interestingly, a similar reduction in ER lumen diameter results from depleting Climp63, where the ER lumen is also decreased to a diameter of 30 nm (Shibata et al., 2010), potentially suggesting that NOMO might maintain this smaller diameter of 30 nm.

**Figure 6.**
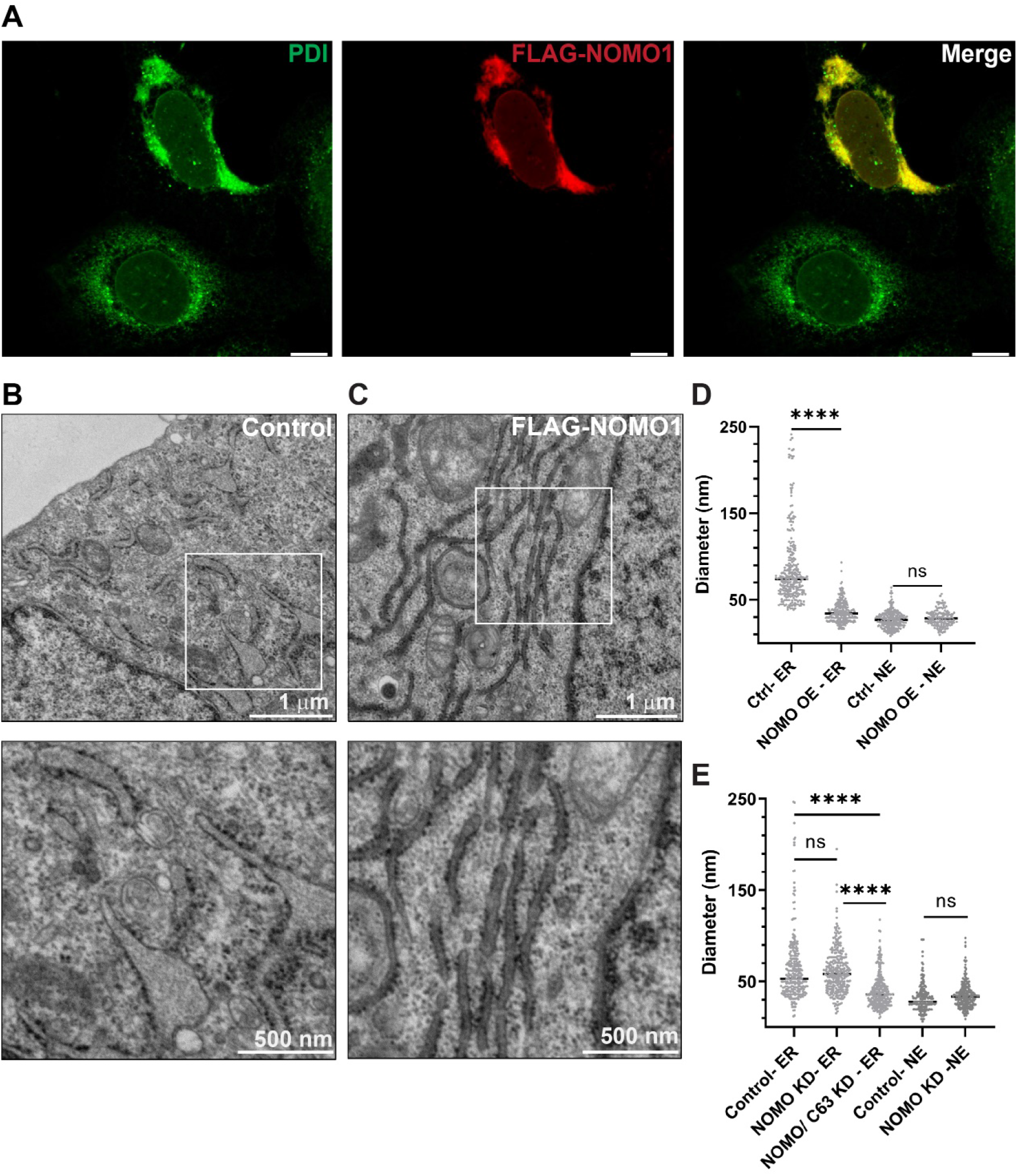
NOMO1 restricts the lumen of the ER. A. Localization of FLAG-NOMO1 (left panel) with ER marker, PDI (middle panel) in U2OS cells. B. HeLa cells transfected with empty vector pcDNA3 as a control. White box in top panel identifies selected zoomed ER membrane area in bottom panel. C. HeLa cells transfected with FLAG-NOMO1. White box in top panel identifies selected zoomed ER membrane area in bottom panel D. Quantified diameters of ER and NE cross sections from A and B. OE = overexpression. Asterisks indicate P<0.0001, ns= not significant. E. Quantified diameters of ER and NE cross sections of U2OS cells treated with the respective siRNAs. Asterisks indicate P<0.0001, ns = not significant.

If NOMO could be the sole remaining ER sheet spacer, we reasoned that simultaneous depletion of NOMO and Climp63 would result in a synthetic effect. Would it become wider than the ER diameter in a wild type cell? The diameter could alternatively decrease as sheet shaping proteins have been proposed to help keep the opposing sheet membranes from collapsing into each other (Schweitzer et al., 2015). To address this question, we simultaneously depleted U2OS of NOMO and Climp63 and processed the cells for electron microscopy. The ER lumen remained restricted and had an average diameter of 40 nm (Fig. 6E), which was significantly less than the control sample, 63 nm. This result does not fit either hypothesis and instead suggests that ER sheet morphology is not dependent on these two proteins alone.

### NOMO is a rod-shaped dimer

Since the overexpression of NOMO causes a uniform restriction of ER intermembrane spacing, we hypothesized that NOMO may support sheet structure by dimerizing across the sheet membranes to support the luminal diameter as originally proposed for Climp63 (Shibata et al., 2010). To determine if NOMO could oligomerize, NOMO1-FLAG was purified from Expi293F cells and analyzed by size exclusion chromatography (Fig. 7A). NOMO1-FLAG eluted at an apparent mass of about 500 kDa based on elution position, which would correspond to a tetramer of NOMO1. However, the potentially elongated form and the correspondingly large apparent stokes radius of NOMO1 could be contributing to an experimental error.

**Figure 7.**
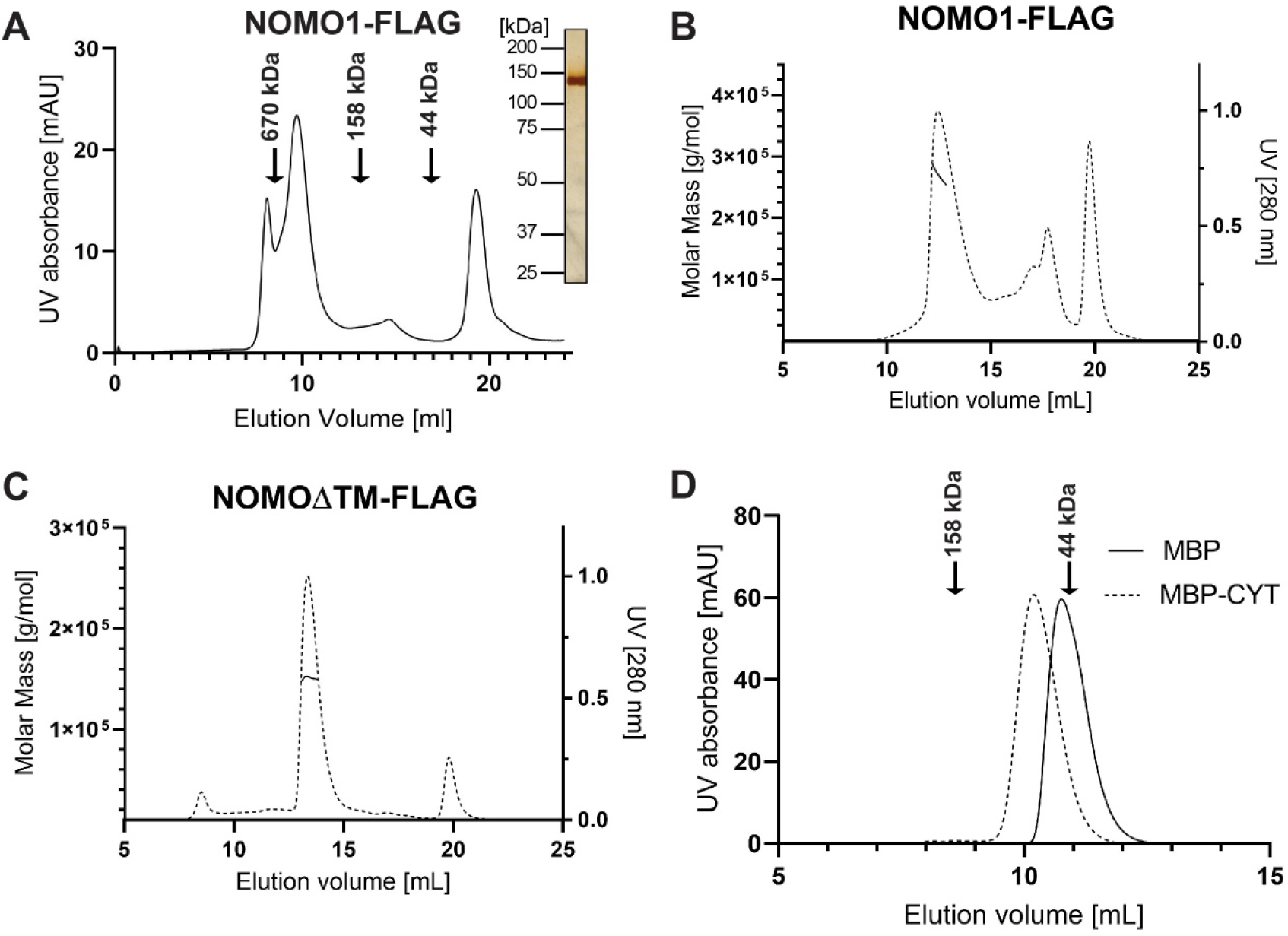
A. Determination of the NOMO1 oligomeric state. A. Elution profile of NOMO1-FLAG on a S200 column. Insert: SDS-PAGE/silver stain of NOMO1-FLAG fraction obtained from preparative SEC. B. SEC-MALS profile of NOMO1-FLAG on a Superose 6 column. Dashed line is the elution profile, solid line is the light scattering profile. C. SEC-MALS profile of NOMOΔTM-FLAG on a Superose 6 column; dashed and solid line as defined in B. D. Dashed line is the elution profile of 2xFLAG-MBP-CYT, solid line is the elution profile of 2xFLAG-MBP.

To accurately determine the oligomeric state and molecular mass of NOMO1-FLAG, we coupled size-exclusion chromatography to multiple angle light scattering (SEC-MALS). The SEC-MALS analysis revealed a molecular mass of 269 kDa and a radius of gyration (R_g_) of about 15 nm (Fig. 7B). This mass would be consistent with a NOMO dimer.

We also performed SEC-MALS analysis with a NOMOΔTM-FLAG construct, lacking both the transmembrane domain and the cytosolic domain, to determine if the dimerization was occurring through the luminal domain. Surprisingly, the analysis showed a mass of 140 kDa from a homogenous peak, revealing that the NOMOΔTM construct is in fact a monomer. Thus, the dimerization is likely occurring through the transmembrane and/or cytosolic domain (Fig. 7C). Furthermore, NOMOΔTM-FLAG had a similar R_g_ (~14 nm) as full-length NOMO1-FLAG. These data argue in favor of NOMO forming a parallel dimer.

To directly test if NOMO is dimerizing via the cytosolic domain (CYT), the CYT domain was fused to maltose binding protein (MBP) to yield 2xFLAG-MBP-CYT. 2xFLAG-MBP and 2xFLAG-MBP-CYT were individually expressed and purified from Expi293F cells and subjected to size exclusion chromatography. While MBP eluted at 58 kDa, MBP-CYT eluted at about 90 kDa based on the elution position. We conclude that despite its small size of 4.8 kDa, the cytosolic tail represents a dimerization domain contributing to NOMO dimerization (Fig. 7D).

### NOMO1 adopts a “beads on a string” morphology

As a first step towards a better structural understanding of NOMO1, we set out to determine the overall architecture of the molecule. NOMO1-FLAG was purified from Expi293F cells and the sample was analyzed by negative-stain EM. 2D class averages were generated using RELION from 7,000 particles. The top 2D class averages from the collected data set feature a flexible, extended rod of about 30 nm (Fig. 8A). A 3D model obtained from these data is 27 nm in length, similar to the ER diameter measured by EM under NOMO overexpression (Fig. 6E and Fig. 8B).

**Figure 8.**
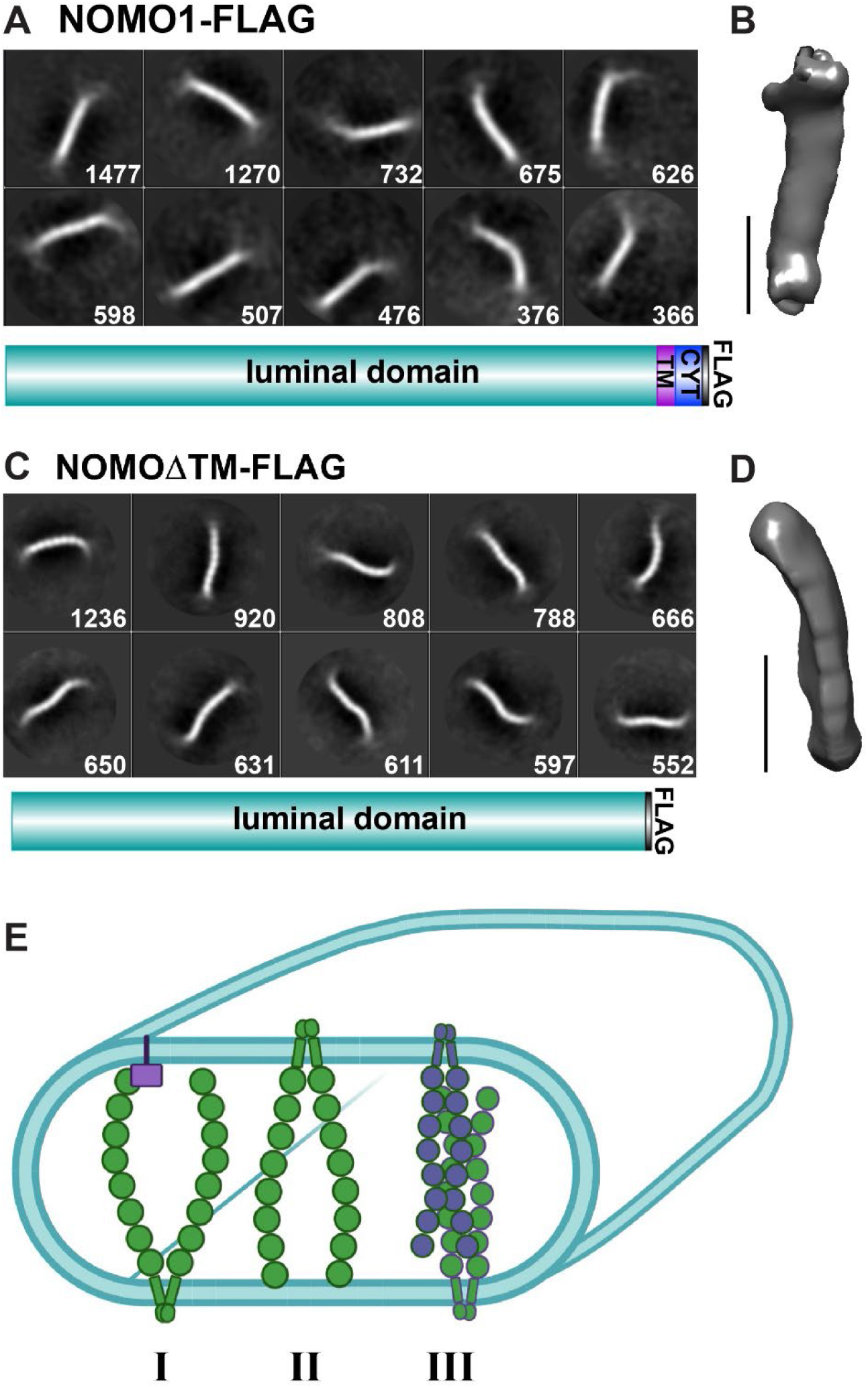
Single particle analysis of NOMO1. A. Top 10 2D class averages of ~7,000 picked negative stain NOMO1-FLAG particles, numbers of particles per class in square. Mask diameter is 40 nm. A construct layout is inserted to clarify protein domains; TM, transmembrane domain, CYT, cytosolic tail. B. 3D reconstruction from A, scale bar is 10 nm. C. Top 10 2D class averages of ~10,000 picked negative stain NOMOΔTM-FLAG particles, number of particles per class in square. Mask diameter is 30 nm. A construct layout is inserted. D. 3D reconstruction from C, scale bar is 10 nm. E. Speculative models for ER sheet imposition by NOMO1. I. Purple block represents an unknown interaction partner on opposite membrane. II. NOMO dimer interacts directly with the opposite membrane. III. NOMO dimers form an antiparallel, low-affinity oligomers.

To determine if a NOMO monomer alone would be similar in length as suggested by the SEC-MALS data and the cytosolic dimerization domain, a negative stain structure was also determined for NOMOΔTM-FLAG since this construct is a monomer. NOMOΔTM-FLAG was purified from Expi293F cells and visualized by negative-stain EM. The 2D classifications were generated using RELION from 9,000 particles and again a flexible and somewhat thinner rod-shaped molecule compared to the full-length protein, consistent with the monomeric nature of this NOMOΔTM construct. Interestingly, the class averages also feature a “beads on a string” morphology with eight discernable globular segments (Fig. 8C), probably accounting for Ig-like domains given the structural homology to bacterial proteins, including BaTIE. The obtained 3D model is about 24 nm in length. In conclusion, NOMO1 is a flexible, rod-shaped parallel dimer featuring a “beads on a string” arrangement of eight consecutive domains, several or all of which may represent Ig-like folds.

## Discussion

In this study, we performed an unbiased proteomics-based experiment to identify abundant, ER-luminal proteins that could serve a function as architectural components of the ER. We identified NOMO1 as an abundant ER constituent of unknown function (Fig. 1), motivating our functional characterization in the context of ER morphology. Notably, NOMO1 and NOMO2 have previously been observed in ER proteomes (Chen, Sans, et al., 2010; Sakai et al., 2009), but remained uncharacterized. NOMO was first described in zebrafish as a nodal signaling regulator (Haffner et al., 2004). The nodal signaling pathway is an embryonic developmental signaling pathway important for cellular differentiation (M. M. Shen, 2007). The ectopic expression of NOMO and Nicalin (NCLN), a NOMO binding partner, leads to cycloptic embryos in zebrafish(Haffner et al., 2004). Transmembrane protein 147 (TMEM147) was later found to form a complex with NOMO and NCLN (Dettmer et al., 2010). NCLN and TMEM147 were recently shown to associate with Sec61 and linked to a role in membrane protein biogenesis (McGilvray et al., 2020). However, the solved structure of this complex did not contain NOMO1, leaving the molecular function of NOMO unresolved.

Our morphological characterization of NOMO depleted cells revealed a drastic rearrangement of the ER network, creating vacuole-like holes in the ER network (Fig. 2A). This phenotype was rescued by overexpression of Atl2 and Climp63. This suggests that the hole phenotype is likely due to an architectural problem since Atl2 and Climp63 provide structural support to the ER, connecting NOMO to the network of known ER shaping proteins.

Ultrastructural analysis of the holes that arise upon NOMO depletion reveal an enrichment of lysosome-like, electron dense structures (Fig. 4A). Consistently, autophagy was altered upon NOMO or Climp63 depletion, as judged by a strong increase in LC3. Of note, NCLN or TMEM147 depletion did not provoke an upregulation of LC3-II (Fig. 5C), and we did not observe rearrangements of the ER network in this experimental context (Fig. S2). Thus, NOMO1 can likely function independently of the NCLN/TMEM147 complex. We did not observe an induction of the UPR in NOMO depleted cells (Fig. 5B), arguing against a critical function for membrane protein biogenesis. However, we cannot formally exclude subtle folding defects that would not amount to a UPR induction. Another possibility is that NOMO could additionally serve as a sheet anchor for the NCLN/TMEM147/Sec61 complex to recruit the process of biogenesis of certain polytopic proteins to flat regions of the membrane.

Regardless, our observation of autophagy dysregulation upon NOMO depletion stresses the relationship of form and function of the ER. Besides imposing a distinct shape on sub-compartments of the ER, ER shaping proteins may additionally be important for defining distinct identities of these compartments. It is interesting to note that while Atl2 depletion results in LAMP1 positive compartments but not an LC3-II increase, Climp63 depletion does not provoke enlarged lysosomes but does result in an LC3-II increase. On the other hand, NOMO depletion causes both a robust increase in LC3 levels and LAMP1-positive compartments (Fig. 5). It will therefore be interesting to closely scrutinize the relationship between autophagy and these membrane-shaping proteins in the future.

Overexpression of NOMO1 resulted in a restriction of the ER luminal diameter to about 30 nm (Fig. 6E). This was particularly interesting because Climp63 depletion results in a decrease of the ER lumen to 30 nm (B. Shen et al., 2019; Shibata et al., 2010), implying that NOMO may be amongst the remaining sheet-shaping proteins responsible for this smaller diameter of 30 nm. Our structural analysis revealed that NOMO1 is an extended, flexible rod of about 27 nm in length, which is similar to the diameter that NOMO1 overexpression imposes on the ER lumen. We speculate that the flexibility of NOMO1 revealed by the negative stain particles may be a structural feature to prevent an overly rigid property of ER sheets. Climp63 had been proposed to be a stable coiled coil (Vedrenne et al., 2005). More recently, calumenin-1 was recently shown to regulate Climp63’s distribution across ER sheets (B. Shen et al., 2019), allowing the ER to adapt and respond to physiological demands that require different distributions of sheets versus tubules.

The NOMOΔTM model revealed that NOMO1 features eight discernable domains that are arranged as “beads-on-a-string” domains (Fig. 8B), reminiscent of the POM152 structure (Upla et al., 2017). Considering that Ig domains can have high structural similarity without significant sequence homology (Berardi et al., 1999), and predicted structural similarity to the bacterial Ig-fold proteins including BaTIE (Fig. S1C), our interpretation is that each of these segments correspond to one Ig fold domain, consistent with the secondary structure prediction showing a high beta sheet content for nearly the entire sequence of NOMO1 (Fig. S1B). Interestingly, structurally related pili proteins in bacteria can dissipate mechanical forces by acting as molecular shock absorbers (Echelman et al., 2016). Thus, it will be interesting to test if NOMO fulfills a similar function in the ER, and to explore possible links to the cytoskeleton.

While our structure-function analysis in concert with light scattering experiments support a model of NOMO1 forming parallel dimers (Fig. 8E), we were not able to unambiguously observe this oligomeric state by negative stain EM, although the dimeric full-length monomer did appear to be thicker than the NOMOΔTM construct. We attribute this problem to the dimerization occurring through the small cytosolic tail, providing for flexibility between the luminal and transmembrane domains.

How can we reconcile the dimensions of NOMO1 with our proposed role as a sheet-shaping protein? We consider three models to relate the dimensions of the NOMO1 rod to the intermembrane spacing observed upon NOMO1 overexpression. First, another, yet unidentified protein interacts with the distal luminal end of the NOMO1 dimer at the opposite membrane (Fig. 8E, I). Second, the distal luminal end of the rod-shaped molecule interacts with the membrane itself (Fig. 8E, II). Third, NOMO1 dimers form antiparallel oligomers of weak affinity (Fig. 8E, III) such that these interactions are not necessarily captured by SEC-MALS analysis in conjunction with size exclusion chromatography. Indeed, a number of distinct oligomeric states of Climp63 were recently observed by analytical ultracentrifugation (J. Zhao & Hu, 2020). If NOMO or Climp63 require an interaction partner to induce their sheet shaping functions (Fig. 8E, I), then overexpressing NOMO or Climp63 would not necessarily cause a striking constriction of the ER intermembrane spacing as the quantity of the interaction partner could be a limiting component. The direct membrane interaction model (Fig. 8E, II) or antiparallel oligomers model (Fig. 8E, III) do not rely on the presence of an interaction partner and could more readily explain the observed correlation between NOMO1 expression levels and sheet formation. Clearly, additional experiments will be required to test these models in the future.

In conclusion, we identified a critical role for NOMO1 in sustaining the morphology of the ER. We propose a dynamic model where both the molecules responsible for membrane spacing and the interactions between them or their interaction partners are highly dynamic. This could be achieved by the inherent flexibility of membrane spacing proteins as exemplified by NOMO1, as well as low to moderate affinity interactions with binding partners at the opposite membrane. In line with this model, homotypic Climp63 interactions appear to be weak (J. Zhao & Hu, 2020). A dynamic model relying both on avidity of multiple weak interactions and inherent flexibility would ensure that ER spacers do not form an impediment for the secretion of bulky cargo (e.g. procollagen with 300-450 nm in length (Malhotra & Erlmann, 2015)), and allow for rapid adjustments of the ER morphology in response to physiological demand.

## Materials and Methods

### Tissue Culture and Stable Cell Line Generation

U2OS and HeLa cells from ATCC were maintained at 37⁰C, 5% CO2 and regularly passaged in DMEM media supplemented with 10% (vol/vol) Fetal Bovine Serum (Gibco) and 1% (vol/vol) Penicillin/Streptomycin (Gibco). Expi293F cells were maintained at 37⁰C, 8% CO2 in Expi293F Expression Media and passaged to maintain a density of less than 8 million cells per mL.

U2OS and HeLa cells were transfected with plasmids using X-tremeGene9 or Fugene-6, according to the manufacturer’s protocol, 24 hours before fixing with 4% paraformaldehyde in phosphate buffered solution (PBS). For rescue assays, U2OS cells were co-transfected with the DNA plasmid and siRNA using Lipofectamine 2000 for 48 hours.

For siRNA transfections, RNAi Lipofectamine was used to transfect U2OS and HeLa cells. siRNA was used at a final sample concentration of 50 nM. A double dose protocol was followed for NOMO and Climp63 depletion where the cells were transfected with siRNA on the first day, transfected again with siRNA 24 hours later, and fixed with 4% (vol/vol) paraformaldehyde in PBS 48 hours after the second transfection.

NOMO and Climp63 were depleted with ON-TARGETplus SmartPools from Dharmacon. Atlastin2 was depleted using the siRNA as in (Pawar, Ungricht, Tiefenboeck, Leroux, & Kutay, 2017).

### APEX2 and Mass spectrometry

ER-APEX2 was transfected into 2 × 10 cm plates of HeLa cells using XtremeGene-9 and expressed overnight. 16-18 hr later, cells were incubated with 500 μM biotin-phenol for 30 min and then treated with 1 mM hydrogen peroxide, from a freshly diluted 100 mM stock, for 1 min before being quenched with 2× quenching buffer. 2× quenching buffer contained 50 mg Trolox and 80 mg sodium ascorbate in 20 mL of phosphate buffered solution (PBS). Cells were rinsed with 1× quenching buffer twice and once with PBS. One control plate was not treated with hydrogen peroxide, but was still rinsed with 1×quenching buffer and PBS. 0.05% Trypsin was then added to the cells for collection into a microfuge tube. Cell samples were spun down at 800 g, 3 min, 4⁰C, rinsed once with PBS, spun down again at 0.8 g, 3 min, 4⁰C, then lysed in an SDS buffer, before quantifying protein concentrated with a BCA Assay (Thermo Fisher). The original protocol can be found in (Hung et al., 2016) Equal amount of lysate samples were incubated with 30 μL streptavidin resin for 3 hours. The beads were washed 3 times and then eluted using 2 × Laemmli Sample Buffer (Bio-Rad). The elution was subjected to SDS PAGE to run the sample into the lane. The lane was then excised into two to three bands and submitted for mass spectrometry analysis.

Mass spectrometry samples were analyzed by the Mass Spectrometry (MS) & Proteomics Resource of the W.M. Keck Foundation Biotechnology Resource Laboratory located at the Yale School of Medicine. An LTQ-Orbitrap XL was used (Thermo Scientific).

### Immunofluorescence

Imaged cells were fixed in 4% (vol/vol) paraformaldehyde/PBS for 15 minutes and permeabilized with 0.1% Triton X-100/PBS for 10 minutes before blocking with 4% (wt/vol) BSA/PBS for another 10 minutes. Samples were then incubated with primary antibodies diluted to 1:500 in 4% BSA/PBS and secondary antibodies diluted to 1:700 in 4% BSA/PBS for one hour each. Samples were rinsed three times with PBS between and after antibody incubations and mounted onto slides using Flouromount-G (Southern Biotech).

For samples where the LAMP1 antibody was used, a gentle permeabilization method was followed. After being fixed in 4% (vol/vol) paraformaldehyde/PBS for 10 min, cells were gently permeabilized with a solution of 0.05% (wt/vol) saponin and 0.05% (vol/vol) NP-40/ PBS for 3 min. The cells were then rinsed with 0.05% saponin/PBS and incubated with primary and secondary antibodies respectively diluted in 0.05% saponin, 1% BSA/ PBS. Samples were then rinsed with PBS and mounted onto slides using Flouromount-G.

Quantification on ImageJ of band intensity was done by converting the immunoblot image to an 8-bit image and creating a binary image to highlight and convert the relevant bands to pixels. The pixels were then measured with the “Analyze Particles” tool. “Show: Results” was selected to label the bands in a binary image with relevant pixel quantification.

### Antibodies

The antibodies used include the following: Protein disulfide isomerase (PDI), Abcam, ab2792. BiP, Abcam, ab21685. Actin, Abcam, ab8226. Alpha-Tubulin, Sigma, T5168. LAMP1, BioLegend, 328602. Calnexin, Abcam, ab75802. FLAG, Sigma, F1804. LC3, Novus, NB100-2331.

### Transmission Electron Microscopy

The Center for Cellular and Molecular Imaging Electron Microscopy Facility at Yale School of Medicine prepared the samples. Cells were fixed in 2.5% (vol/vol) glutaraldehyde in 0.1 M sodium cacodylate buffer plus 2% (wt/vol) sucrose, pH 7.4, for 30 min at room temperature and 30 min at 4C. After rinsing, cells were scraped in 1% (wt/vol) gelatin and centrifuged in a 2% (wt/vol) agar solution. Chilled cell blocks were processed with osmium and thiocarbohydrazide-omsium liganding as previously described (West et al, 2010). Samples were incubated overnight at 60⁰C for polymerization. The blocks were then cut into 60-nm sections using a Leica UltraCut UC7 and stained with 2% (wt/vol) uranyl acetate and lead citrate on Formavar/carbon-coated grids. Samples were imaged using a FEI Tecnai Biotwin at 80 Kv, equipped with a Morada CCD and iTEM (Olympus) software for image acquisition.

### Cloning, Expression and Purification of NOMO constructs

The following constructs were cloned using Gibson assembly from Dharmacon plasmids containing the original gene into a pcDNA3.1+ vector with a C-terminal FLAG tag: NOMO1-FLAG, FLAG-CLIMP63, ATL2-FLAG. NOMOΔTM-FLAG was subcloned from NOMO1-FLAG using Gibson assembly to include only residues 1-1160. FLAG-NOMO1 was closed using the Dharmacon cDNA to PCR residues 33-1226 into a pcDNA3.1+ vector with an N-terminal MHC I signal sequence follow by a FLAG tag.

Expi293F cells were transfected with the construct of interest using the ExpiFectamine 293 Transfection Kit (Gibco) following the manufacturer’s protocol for a 50 mL culture. Cells were harvested 72 hours post transfection and frozen at −80⁰C. Cell pellets were thawed on ice and lysed in Buffer A (50 mM MES, 100 mM NaCl, 50 mM KCl, 5 mM CaCl_2_), 5% glycerol, and 1% DDM for 1 hour at 4⁰C. Afterwards, samples were spun for 30 minutes at 20,000 g, 4⁰C. The supernatant was incubated with anti-FLAG M2 beads (Sigma) overnight and then loaded into a gravity column for washing before incubating with elution buffer containing 5 μM FLAG peptide for 30 min. The elution was then concentrated to 0.5 mL and subjected to size exclusion chromatography in an S200 or S75 column (GE healthcare). 0.05% DDM was added to Buffer A for full length NOMO and 0.005% DDM for NOMOΔTM, 2XFLAG-MBP-CYT, and 2XFLAG-MBP.

### Size exclusion chromatography linked to multi-angle light scattering (SEC-MALS)

Multiangle laser light-scattering experiments were performed at room temperature in a 50 mM MES pH (6.0), 150 mM KCl, 5 mM MgCl2, 5 mM CaCl2, 2% (vol/vol) glycerol, 0.05% (wt/vol) DDM buffer. Light-scattering data were collected using a Dawn Heleos-II spectrometer (Wyatt Technology) coupled to an Opti-lab T-rEX (Wyatt Technologies) interferometric refractometer. Samples (500 uL) were injected and run over a Superose 6 Increase 10/300 GL column (GE Healthcare) at a flow rate of 0.5 ml/min. Light scattering (690 nm laser), UV absorbance (280 nm), and refractive index were recorded simultaneously during the SEC run. Before sample runs, the system was calibrated and normalized using the isotropic protein standard, monomeric bovine serum albumin. Data were processed in ASTRA software as previously described (Wyatt, 1993).

### Single Particle Electron Microscopy

3.5 μL of purified NOMO1-FLAG or NOMOΔTM-FLAG were negatively stained using 2% uranyl acetate solution on carbon film, 400 mesh copper grids that were glow discharged. Grids were imaged on a FEI Talos L120C Electron Microscope (Thermo Fisher Scientific) at 120 kV. Micrographs were captured at a magnification of 73,000x. 82 and 61 micrographs were taken for NOMO and NOMO deltaTM, respectively. TIFF files were cropped to 4096×4096 pixels and converted to MRC format using the EMAN2 v2.3 (Tang et al., 2007) *eproc2d* program. 2D clasifications and 3D reconstructions were produced using RELION v3.08 (Scheres, 2012) with manually picked particles. CTF estimation was performed using using CTFFIND-4.1 with box sizes of 512 and 352 pixels for NOMO and NOMO delta TM, respectively. Particles were extracted, then downscaled four-fold for 2D class averages. Selected 2D classes used for 3D reconstruction are shown in Fig. 8. Final 3D volumes were generated by applying masks generated from initial models and auto-refinement in RELION.

## Acknowledgements

This work is supported by National Institutes of Health grants R01GM114401 and GM126835 (to C. Schlieker), CMB TG T32GM007223 (to C. Amaya), and the National Science Foundation Graduate Research Fellowship DGE1752134 (to C. Amaya). We thank Morven Graham for skilled help with EM, the MS & Proteomics Resource at Yale, and the Schlieker laboratory for critical reading of the manuscript.

This work is supported by National Institutes of Health grants R01GM114401 and GM126835 (to C. Schlieker), CMB TG T32GM007223 (to C. Amaya), and the National Science Foundation Graduate Research Fellowship DGE1752134 (to C. Amaya). We thank Morven Graham for skilled help with EM, the MS & Proteomics Resource at Yale, and the Schlieker laboratory for critical reading of the manuscript.

## Materials and Methods

### Tissue Culture and Stable Cell Line Generation

NOMO and Climp63 were depleted with ON-TARGETplus SmartPools from Dharmacon. Atlastin2 was depleted using the siRNA as in (Pawar et al., 2017).

## Figures and Figure Legends

**Supplementary Figure 1.**
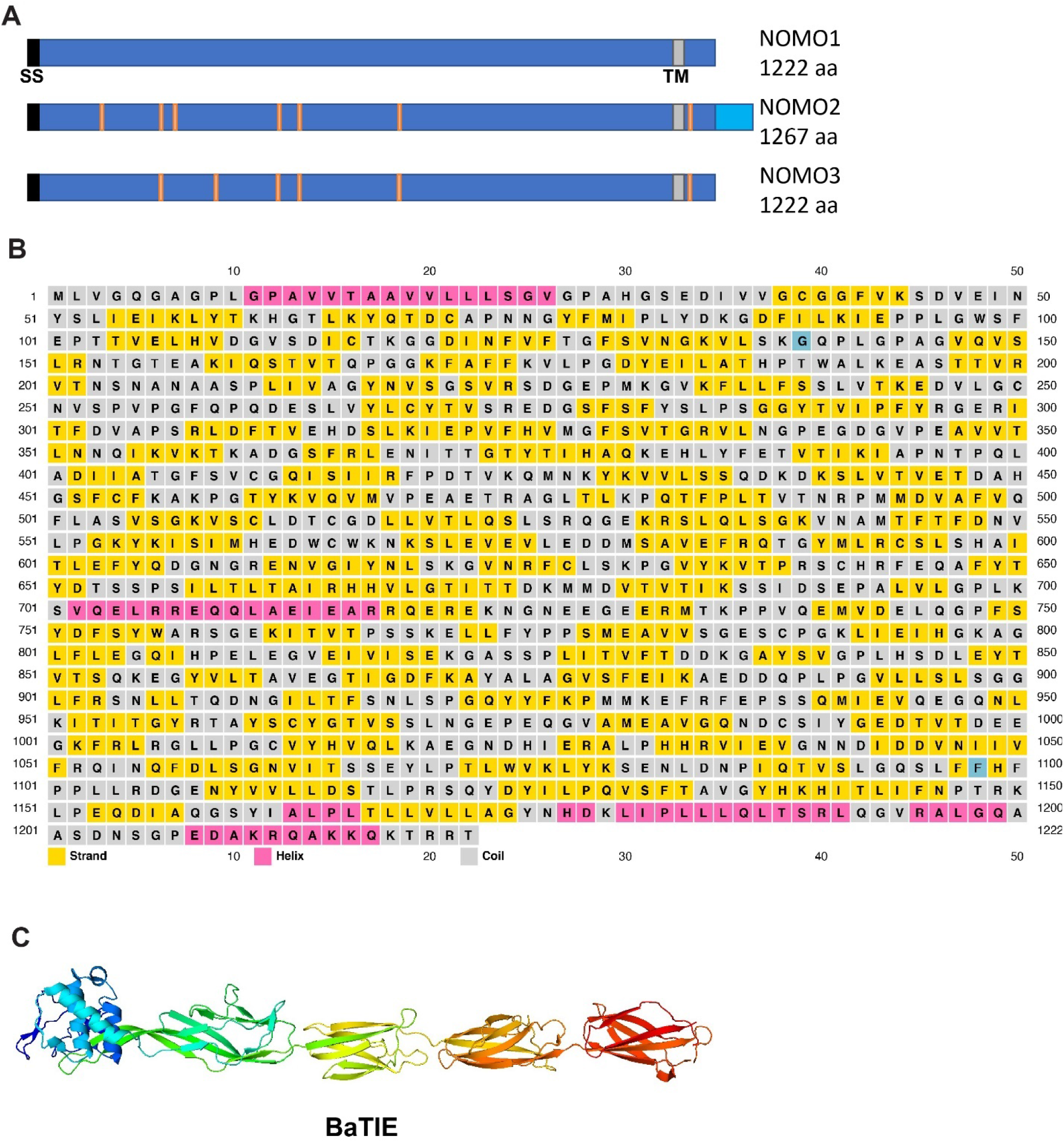
Domain organization of NOMO1. A. NOMO1, NOMO2, and NOMO3 isoforms. Orange denotes single amino acid differences between isoforms. SS= signal sequence. B. NOMO1 secondary structural prediction from PSIPRED. C. BaTIE structure, PDB: 6FWV.

**Supplementary Figure 2.**
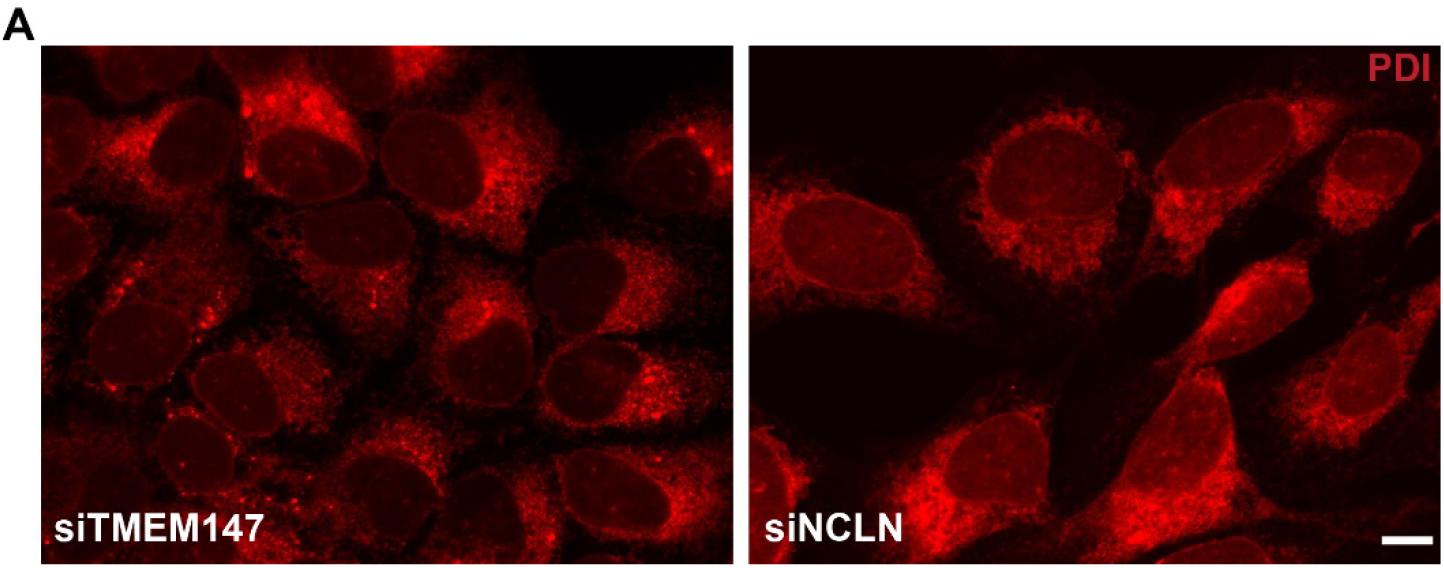
Depletion of NOMO interaction partners do not cause ER morphology disruptions. A. U2OS cells were treated with the denoted siRNAs for 48 hrs and imaged via immunofluorescence. Scale bar is 10 μm.

